# A high-throughput approach to predict A-to-I effects on RNA structure indicates a change of double-stranded content in non-coding RNAs

**DOI:** 10.1101/2022.05.30.494057

**Authors:** Riccardo Delli Ponti, Laura Broglia, Andrea Vandelli, Alexandros Armaos, Marc Torrent Burgas, Natalia Sanchez de Groot, Gian Gaetano Tartaglia

## Abstract

RNA molecules undergo a number of chemical modifications whose effects can alter their structure and molecular interactions. Previous studies have shown that RNA editing can impact the formation of ribonucleoprotein complexes and influence the assembly of membrane-less organelles such as stress-granules. For instance, N6-methyladenosine (m6A) enhances SG formation and N1-methyladenosine (m1A) prevents their transition to solid-like aggregates. Yet, very little is known about adenosine to inosine (A-to-I) modification that is very abundant in human cells and not only impacts mRNAs but also non-coding RNAs. Here, we built the CROSSalive predictor of A-to-I effects on RNA structure based on high-throughput in-cell experiments. Our method shows an accuracy of 90% in predicting the single and double-stranded content of transcripts and identifies a general enrichment of double-stranded regions caused by A-to-I in long intergenic non-coding RNAs (lincRNAs). For the individual cases of NEAT1, NORAD and XIST, we investigated the relationship between A-to-I editing and interactions with RNA-binding proteins using available CLIP data. We found that A-to-I editing is linked to alteration of interaction sites with proteins involved in phase-separation, which suggests that RNP assembly can be influenced by A-to-I. CROSSalive is available at http://service.tartaglialab.com/new_submission/crossalive.

## INTRODUCTION

Although 80% of the human genome is transcribed into RNA, only 2% is translated into proteins (mRNAs), and the rest constitutes the abundant and diverse set of non-coding RNAs (ncRNAs) (1). These RNAs have many important roles such as the regulation of translation, RNA splicing and DNA replication, but also in chromosome structure and immunity (2). This functional diversity is reflected in the wide range of lengths and conformations that ncRNAs can adopt. Accordingly to their size, they can be classified into small ncRNAs (< 200□nt) and long ncRNAs (lncRNA, >200nt). Moreover, this diversity can be furtherly expanded through the addition of post-transcriptional modifications to the nucleotide chain.

As structure confers strength and specificity to molecular interactions, RNA regulation tightly depends on structural properties and their effects on molecular partners recruitment (3). Thus, not only structure determines stability and spatial configuration of RNA molecules but the molecular partners define RNA localization and function. In the case of lncRNAs, their length provides room to incorporate multiple interaction sites providing high valency. As a result, one molecule of RNA can bind several proteins (3, 4). This property is crucial for the formation of ribonucleoprotein (RNP) condensates such as stress granules (SGs) and P-bodies (PBs) where lncRNAs can attract proteins and RNAs (5). Accumulation of proteins and RNAs is a crucial event driving the formation of SGs and PBs via phase separation (4, 6, 7). In biomolecular condensates, RNAs form dynamic interactions with proteins and other RNAs, favoring the exchange of energy and components with the surrounding environment (6). Although all RNAs can be in principle recruited within SGs and PBs (8, 9), some lncRNAs play a special role in this context, such as NORAD, which controls the genomic stability in mammalian cells (10). In order to do so, NORAD can sequester and inhibit Pumilio (PUM) proteins by promoting the formation of phase-separated PUM condensates called NP bodies. Disruption of these condensates leads to genomic instability due to PUM hyperactivity (11, 12). Similarly, the architectural lncRNA NEAT1, has modular domains specifically arranged to bind NONO proteins and form paraspeckle compartments (13). Another interesting example is XIST that is responsible for the X-chromosome inactivation. This is achieved by triggering the formation of large assemblies containing components of Polycomb repressive complexes (PRC1 and PRC2) and many other proteins like CELF, PTBP1 and SPEN (7).

Initially, computational approaches were the only affordable strategies, both technically and timely, to assess RNA structure. However, the lack of experimental data for their training limited their precision (11, 12) until new experimental strategies became available. These approaches can be classified as nuclease cleavage and small molecule-based probing. The former employs nucleases to cut single or double-stranded RNAs (RNases P1, S1 and V1), and the cutting pattern shows the location, at a single nucleotide resolution, of the different strand types (16). The latter employs small chemicals (1M7, DMS, N3-kethoxal and NAI-N3) that bind single-stranded RNA bases and interfere in the reverse transcription step. Then, the pattern of transcription errors (prompt stop or mutations) is used to calculate a structural score for each base (17, 18). Overall, these techniques have allowed achieving comprehensive RNA secondary structure maps of human coding and non-coding transcriptome (19).

These experimental data have fed a second generation of computational tools to predict RNA structure with high precision. In this context, we developed the CROSS algorithm, which is based on four different genome-wide studies that use PARS (RNases S1 and V1) and SHAPE (NAI-N3), together with 3D structural data from X-ray crystallography and NMR (20). CROSS is able to predict the secondary structure propensity of an RNA sequence at singlenucleotide resolution without sequence length restrictions. As in the cases of XIST (21) and SARS-CoV-2 RNA genome (22), CROSS can be used to identify the structural state of sites where RNA-binding proteins (RBPs) interact.

Post-transcriptional chemical modifications can alter RNA structure and properties, and thus modulate its function and contact network, without changing the template genomic DNA (23). N6-methyladenosine (m6A) is one of the most abundant RNA modifications, present in different RNA species such as mRNA, rRNA, tRNA, snRNA and lncRNA (24). However, it is especially abundant in mRNAs where it can be present in many different positions and can promote their recruitment into SGs (25–27). An *in vivo* screening with NAI-N3 (icSHAPE) showed that m6A promotes the transition from double-to single-stranded regions (18). Based on these data, we developed CROSSalive, a method for the prediction of RNA secondary structure *in vivo* with and without m6A modifications (28).

In lncRNAs, the most abundant editing is adenosine to inosine (A-to-I), which renders adenosine similar to guanosine in terms of chemical properties (**Figure 1A-B**) (29). The A-to-I modification is thought to rewire the RNA contact network, both intramolecularly, influencing the secondary structure, and intermolecularly, affecting binding partners and their function (30). Also occurring in mRNAs, the A-to-I modification changes how a codon is read by the ribosome and translated into protein. In both the RNA classes, the ADAR family of enzymes catalyze the hydrolytic deamination reaction that converts adenosine to inosine, where the amino group at position 6 of the purine ring changes to a carbonyl group **(Figure 1A**). In humans, there are three ADAR enzymes (ADAR1, ADAR2 and ADAR3) that catalyze the A-to-I conversion (31). Despite their abundance and effects, A-to-I conversion in lncRNAs has been scarcely studied at the experimental level. A recent computational screening shows that this modification could appear in about 200,000 positions in humans, and 65% of them are located within sites that can affect secondary structure (32, 33). This structural relationship was previously investigated by Solomon and co-workers who used PARS-seq to study the effect of ADAR1 silencing (34). Upon ADAR1 silencing, the authors observed a general decrease in double-stranded content with respect to the single-stranded one. Additionally, Corbet and co-workers reported that ADAR1 depletion triggers SG formation (35). Changes in RNA secondary structure due to A-to-I editing are relevant during viral infection and help the innate immunity to distinguish between viral and human double-stranded RNA (dsRNA) patterns (36).

**Figure 1.**
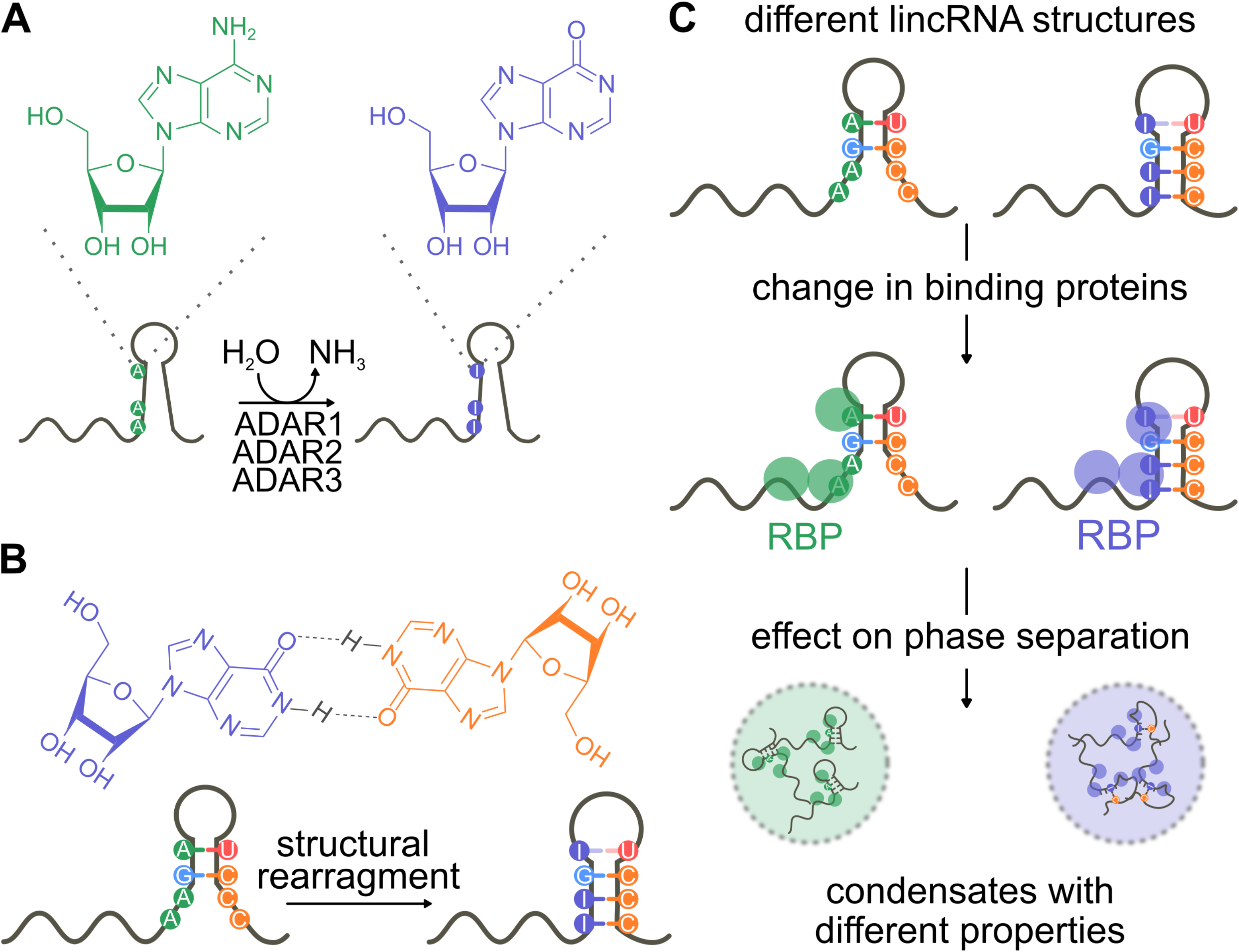
A-to-I editing and its possible implication on condensate formation. **A.** A-to-I editing is catalyzed by adenosine deaminases acting on RNA (ADAR) proteins that convert the adenosine (green) into inosine (purple) by hydrolytic deamination at position C6. **B.** Inosine mimics guanosine and therefore base-pairs favorably with cytosine. Editing causes a structural rearrangement of target RNA. **C.** Modified lincRNAs, due to the altered structure, attract different RNA binding proteins (RBPs) compared to unedited RNAs. We predict that A-to-I editing mediates the recruitment of distinct RBPs and could have an impact on the formation of condensates such as SGs and PBs in the cell.

Here we used Solomon’s PARS-seq data (34) to introduce a new development of CROSSalive that predicts the effects of A-to-I modifications on RNA structure. Specifically, our method calculates the structural profile of a given RNA sequence (single- or double-stranded state) at single-nucleotide resolution in presence (ADAR+) and absence (ADAR-) of ADAR. We employed this version of CROSSalive, available at http://service.tartaglialab.com/new_submission/crossalive, to study the effect of A-to-I modifications on lncRNAs **(Figure 1C)**. Using eCLIP data (37) and *cat*RAPID predictions of protein-RNA interactions (38–40) we investigated the occurrence of A-to-I editing in correspondence of RBPs interactions with three long intergenic non-coding RNAs (lincRNAs), NEAT1, NORAD and XIST. As NORAD, NEAT1 and XIST drive the formation of membrane-less organelles, and our predictions indicate that A-to-I editing alters interaction with proteins prone to phase separation, we hypothesize that A-to-I can have an impact on their macromolecular assembly (**Figure 1C**).

## EXPERIMENTAL PROCEDURES

### PARS data

We processed the read counts from the original PARS experiments on ADAR+ and ADAR-(34) to select nucleotides with a high-propensity to be in single- or double-stranded conformation. In PARS experiments, purified poly-A + RNA samples from control HepG2 cells (ADAR+) and ADAR knockdown (ADAR-) HepG2 cells were extracted and *in vitro* probed with S1 nuclease (S1) and RNAse V1 (V1) that cleave RNA at single-stranded and double-stranded regions, respectively. We used the read counts of S1 and V1 to build a score for each single-nucleotide resolution using the formula Score=log(V1/S1).

### Training set

To build the training set, we followed the approach developed for CROSS (20). Briefly, using the ranking provided by the log(V1/S1) score, we assessed the structural state of each nucleotide (**Figure 2A**) selecting 5 maxima (double-stranded state) and 5 minima (singlestranded state) per transcript. Upon removal of transcripts with insufficient PARS coverage (zeroes for >40% of nucleotides; 3000 sequences passing the filter) and sequence redundancy, we collected a total of 20000 fragments centered around ‘top’ (double-stranded) and ‘bottom’ (single-stranded) nucleotides. Each fragment has a fixed length of 13nt (6 nucleotides on the left and 6 nucleotides on the right side of the central nucleotide). The sequence redundancy of fragments was removed using CD-HIT with threshold at 90% (41). The one-hot encoding (keras in-built function), was employed to convert the fragments into arrays of binary integers for the training of our Artificial Neural Network (ANN).

**Figure 2.**
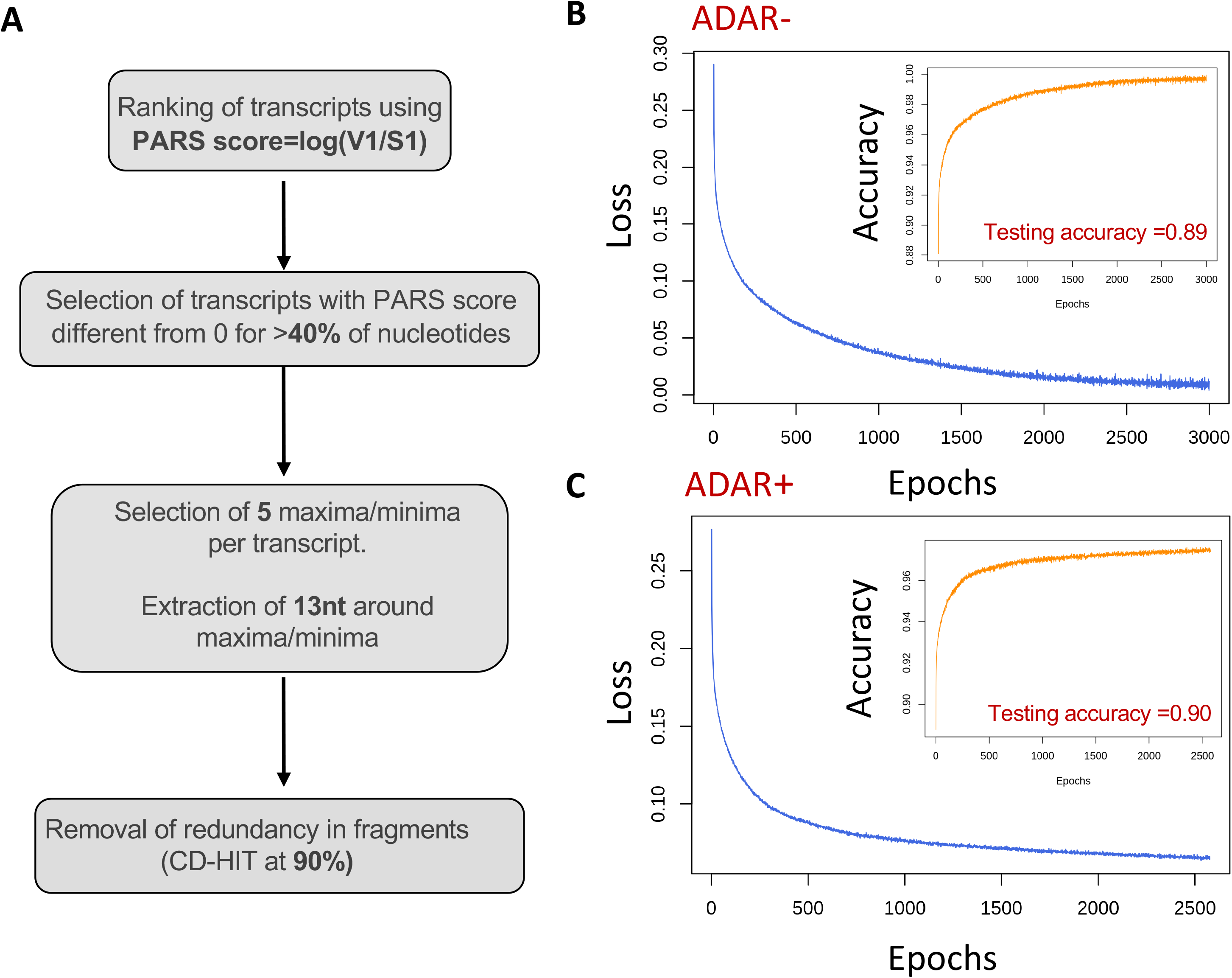
The CROSSalive algorithm. **A.** Workflow for the processing of PARS data, including the selection of high-propensity nucleotides and of the RNA fragments, used for training CROSSalive (log(S1) and log(V1) are the read counts measured upon S1 nuclease (S1) and RNAse V1 (V1) cleaving; **Experimental Procedures**). **B.** Training statistics of ADAR-model including loss (blue line) and training accuracies (orange) at each epoch. The accuracy on an independent test set excluded from the training (i.e. 10% of the training set) is reported. **C.** Training statistics of ADAR+ model including loss (blue line) and training accuracies (orange) for each epoch. The accuracy on an independent test set excluded from the training (i.e. 10% of the training set) is reported.

### Sequences

Sequences of RNAs (20000) reported in the original PARS experiments (34), used in the training, were downloaded from UCSC. cDNA sequences of 22’000 mRNA (**Supplementary Figure 1)** and 12’000 long intergenic non-coding RNAs (lincRNAs; **Figure 3**) were downloaded from Ensembl 105 using Biomart. For mRNAs we used only Ensembl canonical isoforms.

**Figure 3.**
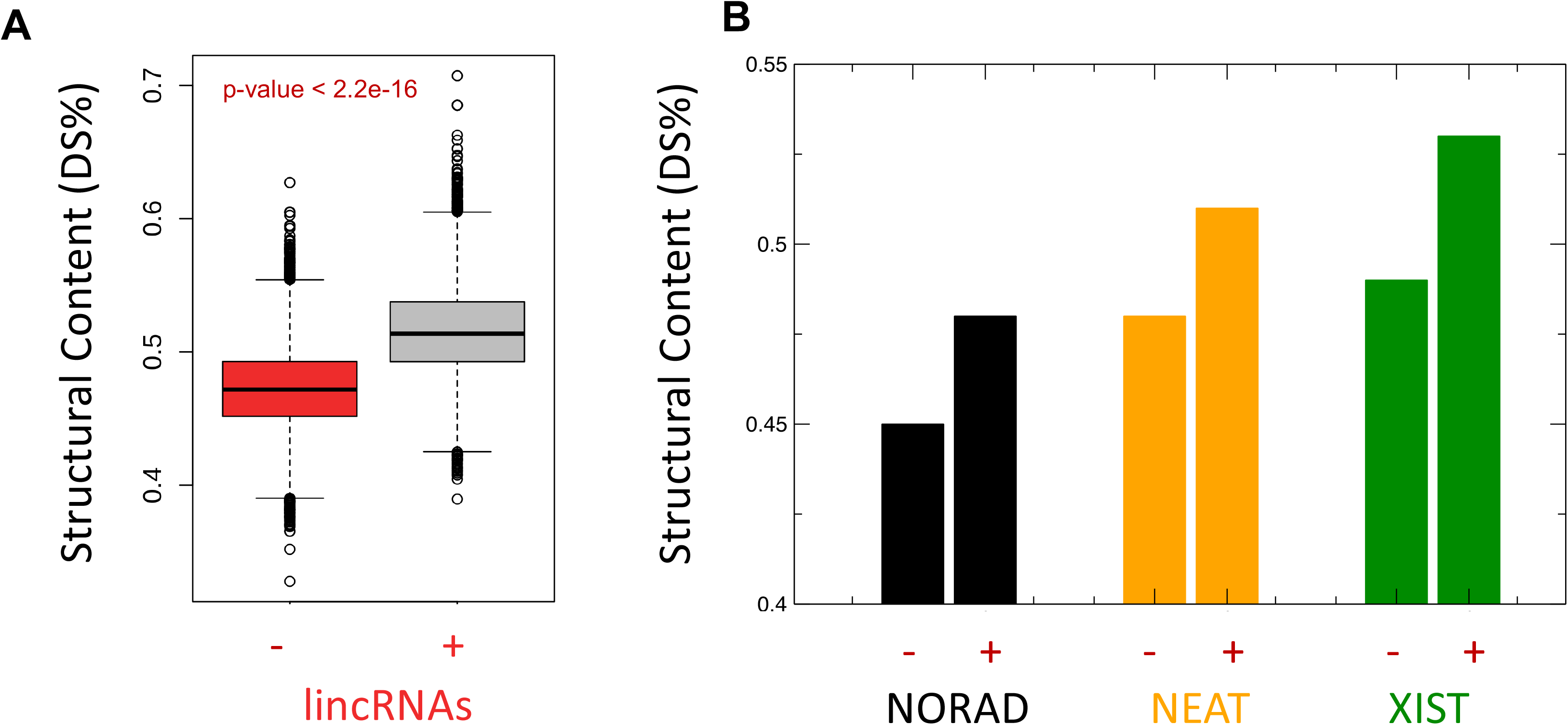
Effects of A-to-I on the structure of long non-coding RNAs. **A.** Boxplots showing the distribution of structural content (i.e. DS% or double-stranded nucleotides) predicted in absence (“-” in red) and presence of ADAR (“+” in gray) for long intergenic non-coding RNAs (lincRNAs). Significance is reported (Kolmogorov-Smirnov). **B.** For lncRNAs of interest (NORAD, XIST and NEAT1), we report differences in the structural content considering absence (“-”) and presence of ADAR (“+”).

**Figure 4.**
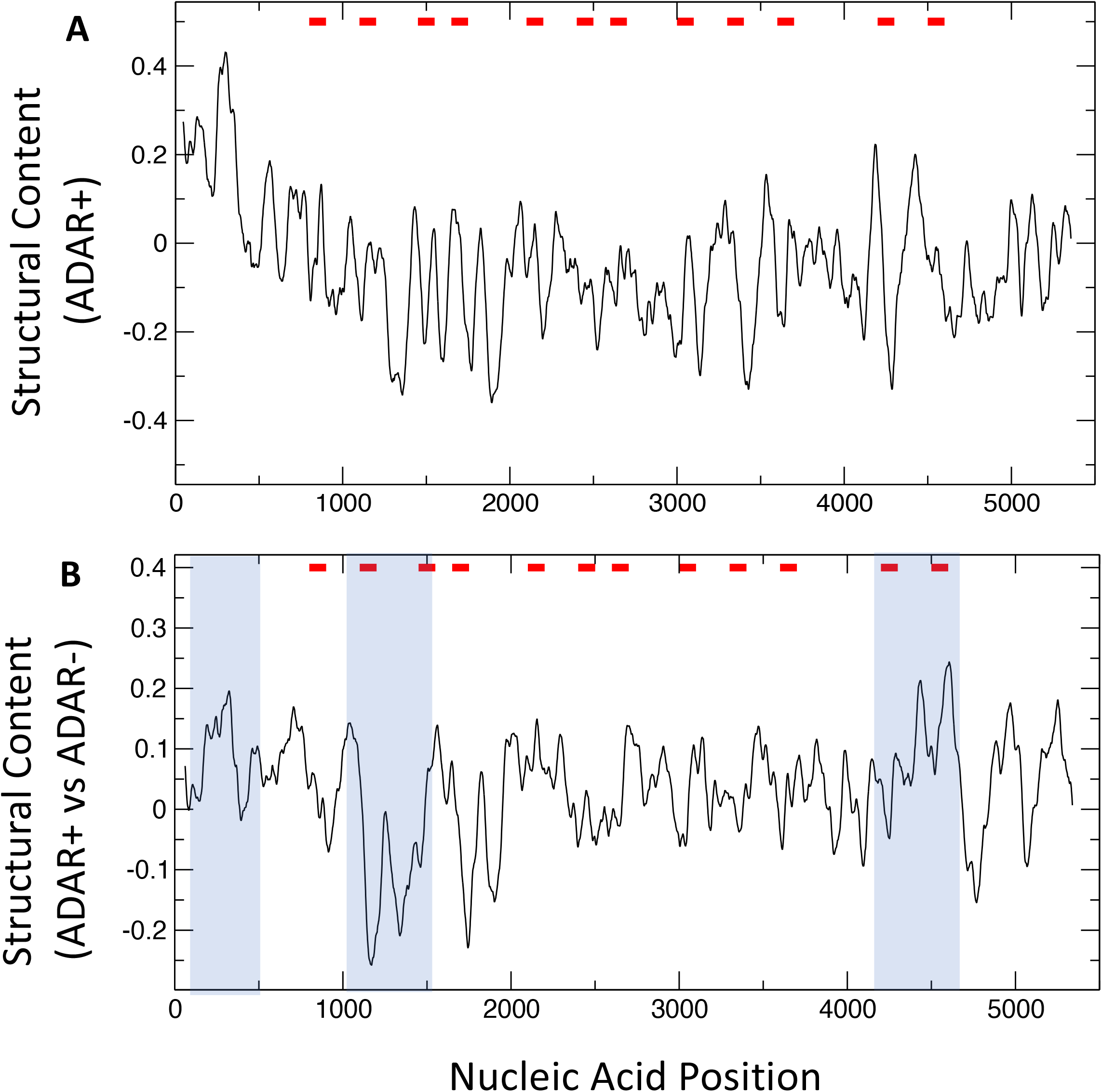
Structural profile of NORAD. **A.** Structural content in presence of ADAR (ADAR+). The 5’ end as well as nucleotides 4100-4650 are enriched in double-stranded regions. **B.** Difference between the structural content in presence (ADAR+) and absence of ADAR (ADAR-). Red bars indicate 12 repeat units (of approximately 300 nt) enriched in the sequence pattern UGURUAUA. Blue boxes indicate regions with the most significant structural changes (i.e. structural peaks and valleys).

### ANN architecture

We used Keras and Tensorflow to train the ANN (42, 43). To keep consistency with our previous algorithm, we used a similar architecture to the one employed for CROSS (20). The architecture is a fully connected Neural Network *(Dense* module in keras), with one hidden layer with 52 internal variables. The layers are based on *relu* and *sigmoid* activation functions, and the training was performed with *adam* optimiser and *binary entropy* loss. The ANN was initially trained for ~3000 epochs, but the final model is trained for 1000 epochs to avoid overfitting (**Figure 2B,C**). 10% of the training set (2000 fragments) was randomly selected and used as an independent test set.

### Performances

We tested the ANNs at each epoch in the training phase by reporting accuracy and loss (**Figure 2B,C**). We used the keras function *train_test_split* to build an independent test set containing 10% (2000 fragments) for ADAR+/- PARS experiments. The networks reached an accuracy of 0.9 on both the independent sets. We tested performances on NEAT1 (>20’000 nt not present in the training set; **Supplementary Information**;) and reported the Area under the ROC curves (AUCs) for both ADAR+/- models using the 10% top (double-stranded) and bottom (single-stranded) nucleotides ranked by log(V1/S1). Both methods achieved an AUC of 0.85 (**Supplementary Figure 2**). Finally, we employed nextPARS (12) experiments on NORAD (**Supplementary Information**) and measured the performances of both ADAR+/- ANNs on the top/bottom 10% nucleotides ranked on nextPARS score, achieving an AUC of 0.77 (**Supplementary Figure 3**). No significant difference exists between PARS and nextPARS protocols, except that nextPARS has higher throughput and sample multiplexing in comparison to PARS (44).

### eCLIP analysis

We used human protein–RNA interactions that were identified through eCLIP experiments in either of the two cell lines, K562 and HepG230 (37). The dataset was downloaded on 18 of September 2020. The dataset contains interactions for 151 unique RBPs: 120 proteins in the K562 cell line and 103 in the HepG2 cell line. We used input-normalized eCLIP data (SM input normalization) with stringent cut-offs [-log10(p value) > 3 and -log2(fold_enrichment) >3]. The data was processed using bioconductor packages for R (https://www.bioconductor.org/) to map genomic locations to Ensembl transcript identifiers.

### *cat*RAPID analysis

To compute protein-RNA interactions, we used catRAPID omics (40). The *cat*RAPID algorithm (38, 39) estimates the binding potential through van der Waals, hydrogen bonding and secondary structure propensities of both protein and RNA sequences allowing identification of binding partners with high confidence. As reported in a recent analysis of about half a million of experimentally validated interactions, the algorithm is able to separate interacting vs non-interacting pairs with an area under the ROC curve of 0.78 (with False Discovery Rate FDR significantly below 0.25 when the Z-score values are > 2) (45) (**Supplementary Information**).

## RESULTS

CROSS (Computational Recognition of Secondary Structure) is a neural network-based machine learning approach trained on experimental data (SHAPE, PARS, NMR/X-Ray, and icSHAPE) able to quickly profile large and complex molecules such as lncRNAs and viral genomes without length restrictions (20). In the original manuscript, CROSS was tested on crystallographic structures (Area Under the ROC curve, AUC 0.72 and positive predictive value PPV 0.74) and on DMS data for murine XIST (AUC 0.75) (20). CROSS was used to profile the HIV genome, reaching an Area Under the ROC curve (AUC) of 0.75 with SHAPE data (20). More recently, CROSS was applied to SARS-CoV-2 (22) and Dengue genomes (46), reaching AUCs of 0.73 and 0.85 with experimental data.

The first version of CROSSalive was trained on icSHAPE data measured in presence and absence of METTL3 (m6A writer) (47). It was also tested on murine XIST (28).

Here, we exploited PARS data on purified poly-A□+□RNA samples from HepG2a experiments by Solomon *et al.* (34) to train a new version of CROSSalive that predicts the effects of A-to-I on the RNA secondary structure profile (**Figure 1**). Briefly, we processed the read counts from the original PARS experiments, both in presence (ADAR+) and absence (ADAR-) of ADAR, to select nucleotides with a high-propensity to be in single- or doublestranded conformation according to the read counts of RNAse V1 (V1) and S1 nuclease (S1) (**Figure 2A**). The information contained in the nucleic acid sequences was converted into arrays of binary integers for the training of a connected neural network, following a similar architecture to previous CROSS algorithms (20) (**Experimental Procedures**). Upon crossvalidation, both ADAR+ and ADAR-networks showed excellent performances, with accuracies of 0.9 in the cross-validation test set (**Figure 2B,C**).

We applied both models (ADAR+ and ADAR-) on human long intergenic non-coding RNAs (lincRNAs) that are a class of lncRNAs that do not overlap exons of either protein-coding or other non-lincRNA types of genes (12’000 transcripts; **Figure 3A**; **Experimental Procedures**). By analyzing the structural content of each transcript (i.e. the percentage of double-stranded nucleotides, DS%), we noticed that there is an increase in structural levels of lincRNAs when predicting their secondary structure using the ADAR+ model. mRNAs also showed a similar trend (**Supplementary Figure 1**), which is in agreement with previous studies in which it was shown that A-to-I modifications promote the formation of secondary structure (32, 34).

With the aim of validating and exploring putative applications of this algorithm, we studied three lncRNAs, NORAD, NEAT1 and XIST. Their analysis revealed that in presence of ADAR, XIST and NEAT1 exhibit a stronger double-stranded content than NORAD (**Figure 3B**).

### NORAD

Systematic investigation of the RNA content of SGs revealed that some lncRNAs such as NORAD are recruited to these condensates (8, 48, 49). NORAD is localized in both the cytoplasm and the nucleus and serves as a platform for PUMILIO (PUM1/2) proteins (50). Although NORAD is not essential for the formation of SGs (49), PUMILIO dependent recruitment of NORAD impacts on SG size and morphology (51). Acting as a PUMILIO sponge, NORAD maintains chromosome stability hampering PUMILIO function (50, 52) and ultimately preventing aberrant mitosis. This is accomplished by the formation of phase-separated condensates *(i.e.* NP bodies) that allow to sequester a high number of PUMILIO proteins. These condensates are generated through the formation of multivalent interactions between NORAD and PUMILIO and also among PUMILIO proteins themselves (6). NORAD contains 12 repeat units (of approximately 300 nt) where the sequence pattern UGURUAUA is repeated (12, 53). Recently it has been shown that NORAD can be methylated at adenosine sites and this would decrease the number of PUMILIO proteins sequestered (54), indicating that RNA modifications can modulate lncRNA activity.

Using CROSSalive we reproduced the secondary structure content, as probed by nextPARS experiments (12), to a remarkable extent, obtaining an AUC of 0.76 for the ADAR+ model and an AUC of 0.77 for the ADAR-model (**Supplementary Image 2; Experimental Procedures**). As the nextPARS experiments were carried out *in vitro* (8), and CROSSalive computes the secondary structure content *in vivo,* the goodness of these performances indicates that NORAD structure does not change significantly in the cell. In presence of ADAR, the most structured regions are the 5’ end of NORAD that interacts with RBMX (52) and nucleotides 4100-4650 that comprise repeats 11 and 12 recognized by PUM1/2 (**Figure 4A**). Notably, NORAD 5’ end region represents the strongest RBMX binding site in the transcriptome (55). While RBMX was first recognized as essential for NORAD function in maintaining genomic integrity (55), it has been recently shown that RBMX is dispensable and that PUM1/2 is necessary for this function (52). It is possible that RBMX and PUM1/2—by interacting with two different NORAD regions—play distinct roles in NORAD-mediated control of genome integrity. Of note, m6A modification induces a structural rearrangement responsible for rendering NORAD less structured and accessible to RBMX binding (56). Since NORAD interactions with RBMX are predicted to occur in highly structured regions according to our predictions, we speculate that m6A and A-to-I modifications could play a role in balancing the amount of RBMX associated with NORAD.

Nucleotides 1000-1500 of NORAD (between repeat 2 and 3) are predicted to be undergoing structural rearrangements in presence of ADAR (blue box in **Figure 4B**). Specifically, our calculations indicate that there is a strong decrease in secondary structure content. Considering RBP interactions detected by eCLIP in this region as (37), we identified FUBP3, IGF2BP2, PUM1, PUM2, TIA1 and TIAL1 that are SG components, which suggests that ADAR could impact NORAD ability to phase-separate. At nucleotides 500-800 and 4300-4650 (repeat 12), we predict that ADAR depletion decreases the secondary structure content. In these regions, the SG component KHDRBS1 binds (57) (blue box in **Figure 4B**). In the 5’ region corresponding to nucleotides 1-500, interactions with SG proteins DDX3X, DDX6, EIF3G, PCBP1 and RPS3 could be potentially affected by changes in the RNA structure (blue box in **Figure 4B**).

### NEAT1

NEAT1 plays a key role in the formation and function of nuclear condensates, namely paraspeckles (58, 59). Paraspeckles—formed through the process of liquid-liquid phase separation—are membraneless subnuclear compartments involved in a variety of cellular processes (60). Paraspeckles not only contribute to gene expression regulation, for instance through sequestration of SFPQ transcriptional factor (61) or by promoting microRNA biogenesis (62), but they also mediate nuclear retention of hyper A-to-I–edited transcripts (63, 64). They are enriched in Drosophila Behavior Human Splicing (DBHS) RBPs such as NONO, SFPQ, and PSPC1, but they also include other RBPs harboring prion like domains *e.g.* FUS, TDP43 and TAF15 (65).

NEAT1 is expressed as two distinct isoforms: a short polyadenylated transcript (NEAT1_1) and longer transcript (NEAT1_2) with a stabilizing triple structure at the RNA 3’end (NEAT1_2) (66). NEAT1_2 is essential for paraspeckle formation, by acting as a scaffold molecule (66). Indeed, assembly of these condensates strictly depends on the transcription of NEAT1_2 and consequently on the proteins that associate to it (58, 67). Genetic analysis revealed that NEAT1_2 contains different functional domains with distinct roles in paraspeckle production (13). Among these domains, the middle region has been demonstrated to recruit the core component NONO, which in turn initiates the paraspeckle assembly (13). NEAT1_2 displays a highly organized spatial arrangement in paraspeckles. Indeed, these condensates form spheroidal structures in which, while the 5’ and 3’ termini of NEAT1_2 are located at the paraspeckles periphery, the middle region is found in the assembly core (59, 68). Super resolution microscopy revealed that NEAT1 attract paraspeckle core componentes *(e.g.* FUS and DBHS family proteins) at the center of the spheroidal structure, while 5’ and 3’ ends of NEAT1_2 and TDBP-43 localize at the paraspeckle shell (59).

NEAT1 has been shown to be chemically modified post-transcriptionally with methylation at cytosines (m5C) and adenosines (m6A) (69, 70). Although the role of RNA modifications on NEAT1 activity needs to be further investigated, recent work has started to shed light on the consequence of m6A modification in NEAT1 and cancer progression (71–73). It has been recently demonstrated that ADAR1 controls NEAT1 stability by editing Alu sequences located in proximity of the AUF1 binding region. The modification decreases the ability of AUF1 to interact with its target (74).

Using CROSSalive we reproduced the secondary structure content of NEAT1 to a remarkable extent (**Experimental Procedures)**, with an overall AUC of 0.85 or more compared to experimental data (34) (**Supplementary Image 3A,B**). As NEAT1 was not used in our training set, this result indicates that CROSSalive reliably predicts structural properties of transcripts.

The 5’ and the 3’ ends as well as nucleotides 5000-8500, 14500-15500 and 17500-19000 are highly conserved, as reported by previous studies (75) and enriched in double-stranded content (**Figure 5A; Supplementary Image 3C**). Comparing ADAR+ and ADAR-profiles of secondary structure content, we found that nucleotides 5000-8500 (corresponding to the highly conserved middle region (59), blue box in **Figure 5B**; see also **Supplementary Image 3C**) and especially nucleotides 7000-8500 undergo major structural rearrangements. This could impact RBP recruitment and ultimately RNP assembly. Indeed, as reported by eCLIP (37) (**Experimental Procedures**), this region involves interactions with proteins that undergo phase-separation including CPEB4, EIF4G2, EWSR1, FUBP3, FUS, HNRNPK, HNRNPM, KHSRP, MATR3, NONO, PCBP1, QKI, SFPQ, TARDBP, TIA1 and TIAL1

**Figure 5.**
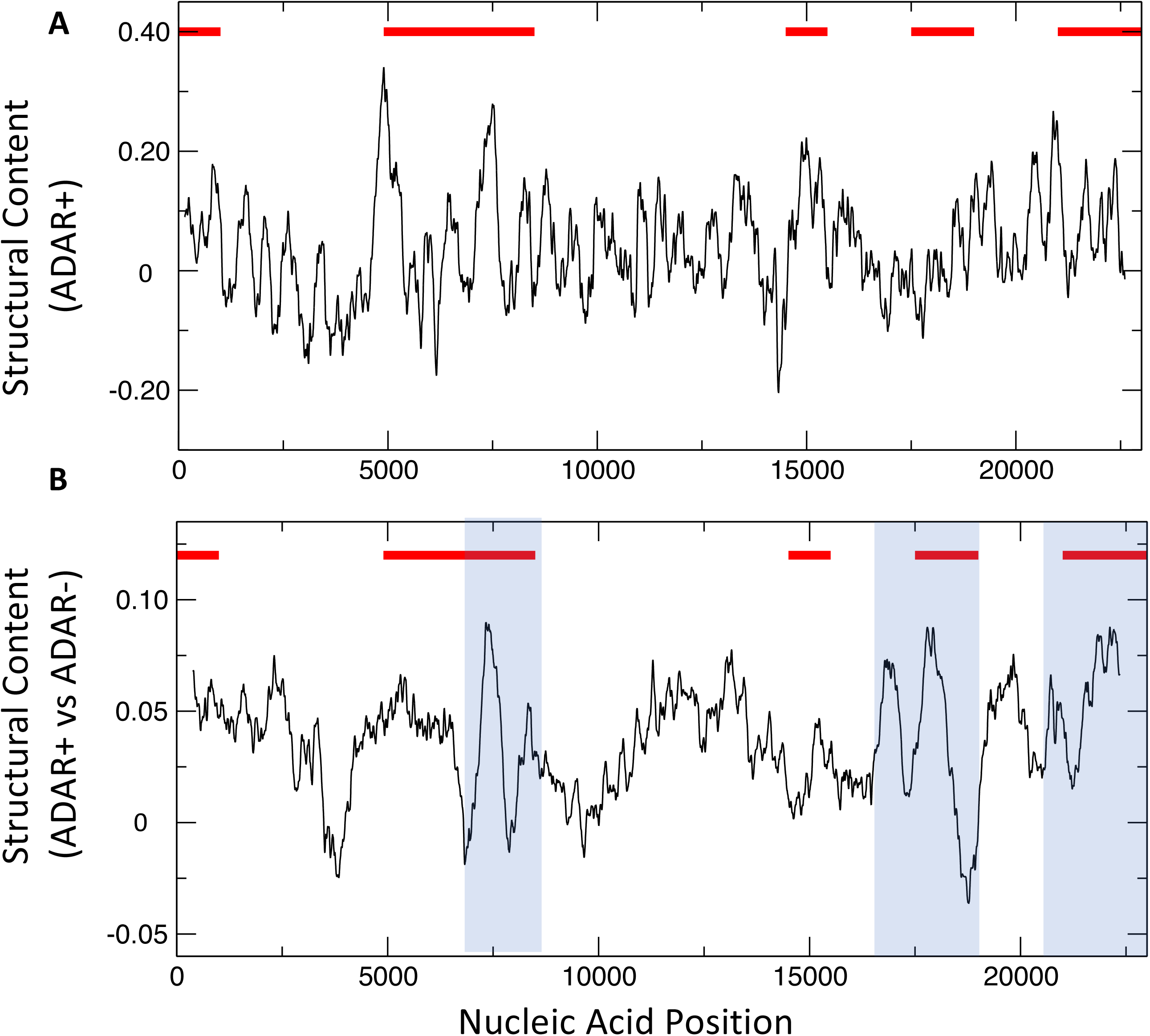
Structural profile of NEAT1. **A.** Structural content in presence of ADAR (ADAR+). The 5’ and the 3’ ends as well as nucleotides 5000-8500, 14500-15500 and 17500-19000 are enriched in double-stranded regions. **B.** Difference between the structural content in presence (ADAR+) and absence of ADAR (ADAR-). Red bars indicate evolutionary conserved regions that are important for paraspeckle structure (59); (**Supplementary Figure 3**). Blue boxes indicate regions with the most significant structural changes (i.e. structural peaks and valleys).

In the 5’ region corresponding to nucleotides 14500-15500 (in the periphery of paraspeckle structure (59); blue box in **Figure 5B**), we predicted a slightly lower decrease in structural content and fewer changes in the binding of phase separating proteins FUBP3, HNRNPK and TIA1 (37). At nucleotides 17500-19000 we predicted an alteration in the structural content upon ADAR depletion but we could not retrieve any protein interactions. In the 5’ tail corresponding to nucleotides 21000-22700, structural changes are accompanied by alteration of the binding sites of DDX6, EIF3G, EIF4G2, EWSR1, FUBP3, FUS, HNRNPK, IGF2BP1, KHDRBS1, NONO, SFPQ, TARDBP, TIA1 and TIAL1 (blue box in **Figure 5B**). In the 3’ region corresponding to the highly conserved region located at nucleotides 1-1000 (in the periphery of the paraspeckle) structure and (59), we found a milder change of structure and few phase-separating proteins affected, including DDX6, EWSR1, FUBP3, FUS, HNRNPM, PCBP2, and TAF15 (37).

Our predictions suggest that A-to-I editing could impact interactions between NEAT1 and proteins crucial for phase separation. The RNA editing could alter the highly ordered spatial distribution of the assembly components within the paraspeckle core and shell.

### XIST

XIST interacts with a plethora of RNA binding proteins (RBPs) involved in gene silencing, RNA modifications and localization. Its interactions with RBPs are mediated by specific RNA regions. XIST harbors six tandem repeats, ‘Rep’, named from A to F, which act as distinct functional domains essential for RBP binding (76–78). According to experimental and computational analysis, XIST Rep A, B and E are more structured than Rep D. XIST Rep A, B and E are crucial for RBP interactions (20, 39).

XIST Rep A at the 5’ region is the most conserved in terms of sequence and copy number and it is probably the most contacted region essential for silencing at the early stages (25, 79). The highly structured Rep A functions as a modular platform for the association of RBPs (*e.g.* SPEN which in turn recruits the SMRT-HDAC3 complex required for gene silencing) and m6A modifications important for XIST functions (80). The cytosine rich Rep B/C modules directly interact with hnRNPK, which subsequently recruit the Polycomb repressive complex 1 (PRC1) (39, 81). The PRC1-mediated ubiquitination of histone H2A is recognized, through the JARID2 cofactor, by Polycomb-repressive complex 2 (PRC2) (82). It has been shown that while the initial recruitment of the Polycomb complexes is mediated by XIST Rep A, the XIST Rep B is crucial in stabilizing the Polycomb complexes and therefore maintaining gene silencing on the inactive X (83). Rep E controls XIST localization, in particular it is important to retain XIST at the inactive X site. Several RBPs have been proved to interact with Rep E, including TDB-43, CELF1, PTBP1 and MATR3 (39, 84, 85). The interaction of these proteins depends on the structural conformation of the Rep E, indicating that it is highly specific (76, 84). It has been hypothesized and proved that Rep E is involved in the formation of phase-separated condensates important for both XIST localization and gene silencing preservation (21, 85). Indeed, it was recently discovered that XIST forms large phase-separated assemblies, similar to stress granules, by progressively attracting proteins (7, 21, 86).

XIST is widely methylated with more than 70 N6-methyladenosine (m6A) residues (79). This modification seems to be crucial for the scaffolding activity of XIST by promoting protein recruitment. For instance, YTHDC1 proteins preferentially recognize m6A residues on XIST (79, 87). It is likely that other RNA modifications also play a crucial role in XIST function and assembly formation. Although ADAR was not identified to bind specifically to XIST in mouse (39), the Lee lab identified two binding partners, Q924C1 and Q9EPU (88). Thus, it is possible that ADAR1 could play an role in XIST activity.

Our calculations indicate that ADAR has the strongest effect on Rep B, C, D and E. Specifically, the double stranded content is significantly reduced upon ADAR depletion (**Figure 6A,B**). The 3’ region of XIST, which does not play any functional role in X chromosome inactivation, is mildly affected by ADAR depletion (**Figure 6B**).

**Figure 6.**
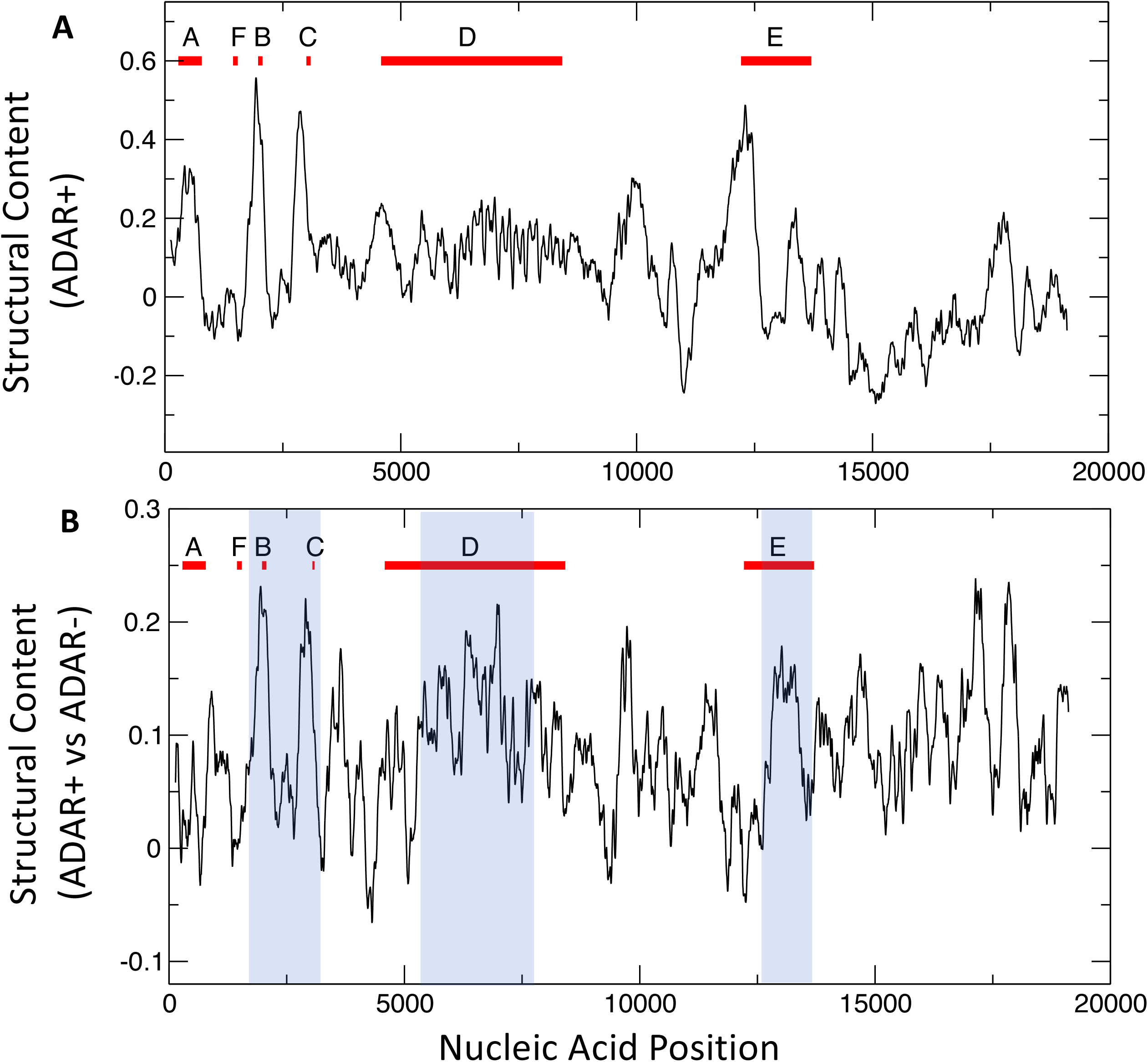
Structural profile of XIST. **A.** Structural content in presence of ADAR (ADAR+). Rep A, B, C, D and E are enriched in double-stranded regions. **B.** Difference between the structural content in presence (ADAR+) and absence of ADAR (ADAR-). Red bars indicate repeat units (78). Blue boxes indicate regions with the most significant structural changes.

Considering RBP interactions in these regions (37), we predict that HNRNPK binding to Rep B may be altered upon A-to-I modifications. A study from the Heard laboratory showed that Rep B is necessary for PRC1 recruitment and the Brockdorff laboratory proved that HNRNPK, which physically interacts with PRC1, is directly involved in RNA binding (89). Importantly, HNRNPK is a SG component (90) and could play a role in XIST phase separation (7, 21). HNRNPK binds to other regions that are altered upon ADAR1 depletion, including Rep D where another SG component binds, HNRNPU (37). Additionally, Rep E interacts with several SG proteins such as MATR3, PTBP1, TARDBP, TIA1 (37).

### A hypothesis on A-to-I effects on protein interactions

Our analysis indicates that structural changes in RNA molecules caused by ADAR could occur in regions contacted by RBPs. However, at present very little is known about the effects of A-to-I editing on RBP interactions.

We used *cat*RAPID (38–40) to investigate the effects of A-to-I modifications on protein binding (**Figure 7**). As the parameters for “I” are not yet integrated in *cat*RAPID, we adopted an approach based on the substitution of “A” with “G” followed by the calculation of RBP interactions. The A-to-I substitution was carried out at random positions within selected regions of NORAD, NEAT1 and XIST corresponding to peaks of ADAR+ profiles.

**Figure 7.**
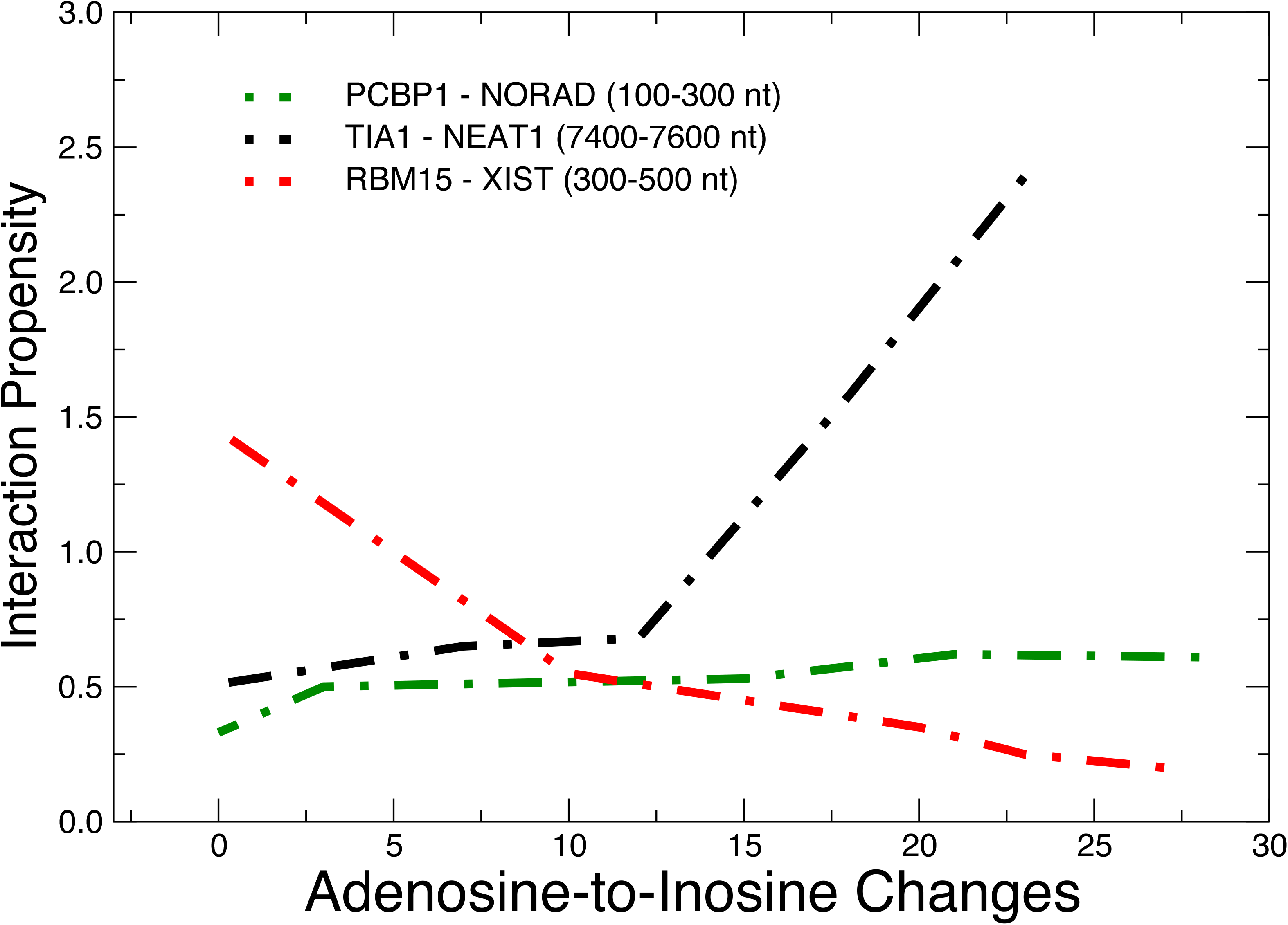
Effects of A-to-I editing on protein binding. *cat*RAPID predictions indicate that PCBP1 only mildly increases its binding propensity for nucleotides 100-300 of NORAD depending on A-to-I modification. By contrast, nucleotides 7400-7600 of NEAT1 associate with TIA1 more tightly when the number of inosines increases. The opposite effect is observed for RBM15 that interacts with Rep A in the 3’ end of XIST. We note that all the interaction propensities are >0, indicating that protein binding is always predicted to occur.

In the case of NORAD, PCBP1 is predicted to only mildly increase its binding propensity for nucleotides 100-300 depending on A-to-I modification (**Figure 7**). This region is the most structured in NORAD (**Figure 4**), and PCBP1 binding might be relevant for phase separation.

As for NEAT1, one of the most structured regions in presence of ADAR (i.e. nucleotides 7400-7600; **Figure 5**) is predicted to associate with TIA1 in a A-to-I dependent manner and, specifically, to interact tightly when the number of inosines increases (**Figure 7**). As the A-to-I editing is observed in adenosines that are in close proximity to the AU-rich of NEAT and TIA1 is a AU-rich binding protein (74), it is tempting to speculate that TIA1 propensity to phase separation might influence NEAT1 ability to form paraspeckles depending on ADAR editing.

The opposite effect is observed for RBM15 that interacts with Rep A at the 3’ end of XIST. In this case, *cat*RAPID predictions indicate that A-to-I modifications decrease the binding ability of RBM15 for nucleotides 300-500 (**Figure 7**). While the structure of this region is only mildly affected by ADAR depletion (**Figure 6**), it is worth mentioning that RBM15 interaction mediates m6A modification of XIST (91), which suggests that A-to-I and m6A can both act in this region.

In summary, our computational analysis indicates that the A-to-I modification can affect RBP interactions in different ways, suggesting that further work in this exciting area is needed.

## CONCLUSIONS

Advancements in next-generation sequencing and mass spectrometry allowed the identification of more than 170 different RNA post-transcriptional modifications (92), and the list is rapidly expanding (93). This finding indicates a complex regulatory network for RNAs and highlights the richness of the whole transcriptome. In an extremely versatile manner, chemical modifications adjust the biological functions of all RNA types (tRNA, rRNA, mRNA, snRNA, lncRNA or viral RNA) by altering their secondary structures, interaction networks, localizations and half-lifes (94).

The contacts that RNAs establish and their spatial locations are crucial for the formation of membrane-less organelles (95). In the mRNA context, several modifications can influence liquid-liquid phase separation. For example, Ries and co-workers showed that mRNAs with multiple m6A sites can act as a multivalent scaffold for the m6A-binding proteins YTHDF (25). he m6A-mRNAs bring YTHDF low-complexity domains closer and trigger phase separation, thus regulating mRNA localization in compartments such as PBs, SGs or neuronal RNA granules, which ultimately affect stability and translation (25). Also, mRNA recruitment into SGs protects transcripts from irreversible aggregation under stress conditions (96). In agreement with this observation, we recently reported that the m1A modification leads to a more efficient recovery of protein synthesis upon heat shock (97). We also found that m1A RNAs and the enzyme involved in this modification (TRMT6/61A) play a role in SG formation (97). These findings indicate that both m6A and m1A can influence transcriptome turnover and translation by modulating the liquid-liquid phase separation process.

The adenosine deamination to inosine (A-to-I) is a very abundant nucleotide modification in animals. Specifically, millions of editing sites have been reported in humans and most of them are located in non-coding regions of the genome (31, 98). Despite its abundance, we still lack an overall understanding of the purpose and functional effects of this modification (31). The development of high-throughput methods is providing a new set of tools to investigate RNA editing.

In this manuscript, we used PARS data to introduce a new development of CROSSalive that predicts how A-to-I modifications affect the RNA structure (34). In lincRNAs, we detected a general enrichment of double-stranded regions, with just a few transcripts having the opposite behavior. For three functional lncRNAs, NEAT1, NORAD and XIST, we studied the relationship between A-to-I editing and protein interactions using available CLIP data (37) and *cat*RAPID (38–40). We found that A-to-I editing is linked to alteration of interactions with several phase separating proteins. This could be a general trend for other lncRNAs (37).

Although protein-RNA interactions are crucial for the formation of membrane-less compartments, emerging evidence has shown that inter- and intra-molecular RNA-RNA interactions are also involved in the RNP condensate formation (51). Relatively to the examples presented here, NORAD tends to lose its structure under stress conditions *(e.g.* arsenite treatment) and becomes more linear. In these conditions NORAD establishes inter-molecular interactions with G/C-rich mRNAs that accumulate in SGs (99). Similarly, the high local NEAT1 concentration at NEAT1 transcription sites allows the establishment of multivalent interactions among NEAT1 molecules, which could shape paraspeckles construction (51, 100, 101). As for XIST, Rep A has been reported to participate in intramolecular interactions forming long repeat duplexes whose architecture is crucial for the assembly of SPEN (101). We cannot exclude that XIST establishes inter-molecular interactions also with other RNAs. Indeed, according to the RISE database, NORAD, NEAT1 and XIST engage in inter-molecular interactions with 18, 150 and 184 RNAs, respectively (102). We envision that alterations of RNA-RNAinteractions—caused by A-to-I editing— undoubtedly have an impact in condensate formation and future research should aim at dissecting the role of lncRNA chemical modifications in RNA-RNA interactions contributing to cellular assemblies.

CROSSalive could be useful to investigate structural changes in contexts different from phase separation. An intrinsic limitation of our method is that it predicts local properties of structure, yet the ability to reproduce experimental data is significantly high (global accuracy of 0.90), which suggests that tertiary structure should play a minor role in the cellular environment, as also previously reported (103). We envisage that an implementation where long-distance interactions are taken into account would make CROSSalive applicable to a broader range of RNA molecules, especially long non-coding RNAs where tertiary interactions are expected to be relevant (104).

In the long term, we aim to expand our suite of algorithms to predict RNA secondary structure changes of other chemical modifications. CROSSalive can now predict structural changes associated with m6A and A-to-I modifications, and we plan to integrate the individual contributions of other modifications as soon as new experimental data will become available.

## ACKNOWLEDGMENTS

The authors would like to thank the ‘RNA initiative’ at IIT and all the members of Tartaglia’s lab at CRG, Sapienza and IIT.

