## Supplementary Information for "A high-throughput approach to predict A-to-I effects on RNA structure indicates a change of double-stranded content in non-coding RNAs"

**CROSSalive analyses**

CROSSalive calculations for the ADAR+ model [are available at http://crg-webservice.s3.amazonaws.com/submissions/2022-05/464934/output/index.html?unlock=42d43cc38c](http://crg-webservice.s3.amazonaws.com/submissions/2022-05/464934/output/index.html?unlock=42d43cc38c) . Similarly, NORAD ADAR- calculations are at <http://crg-webservice.s3.amazonaws.com/submissions/2022-05/464935/output/index.html?unlock=0ee417dec2>, XIST ADAR+ model [is at http://crg-webservice.s3.amazonaws.com/submissions/2022-05/464936/output/index.html?unlock=e516687e57](http://crg-webservice.s3.amazonaws.com/submissions/2022-05/464936/output/index.html?unlock=e516687e57), XIST ADAR- model [is at http://crg-webservice.s3.amazonaws.com/submissions/2022-05/464937/output/index.html?unlock=df60cf2f5d](http://crg-webservice.s3.amazonaws.com/submissions/2022-05/464937/output/index.html?unlock=df60cf2f5d), NEAT1 ADAR+ model [is at http://crg-webservice.s3.amazonaws.com/submissions/2022-05/464938/output/index.html?unlock=9b4a3a3d2b](http://crg-webservice.s3.amazonaws.com/submissions/2022-05/464938/output/index.html?unlock=9b4a3a3d2b) and NEAT1 ADAR- model [is at http://crg-webservice.s3.amazonaws.com/submissions/2022-05/464939/output/index.html?unlock=c91335e350](http://crg-webservice.s3.amazonaws.com/submissions/2022-05/464939/output/index.html?unlock=c91335e350).

***cat*RAPID analysis**

The calculations carried out for this manuscript are are avaiable at <http://service.tartaglialab.com/update_submission/462781/9e0def89fe> (NEAT1), <http://s.tartaglialab.com/update_submission/462881/305aa08ad3> (NORAD) and <http://s.tartaglialab.com/update_submission/463367/15e82ed813> (XIST).

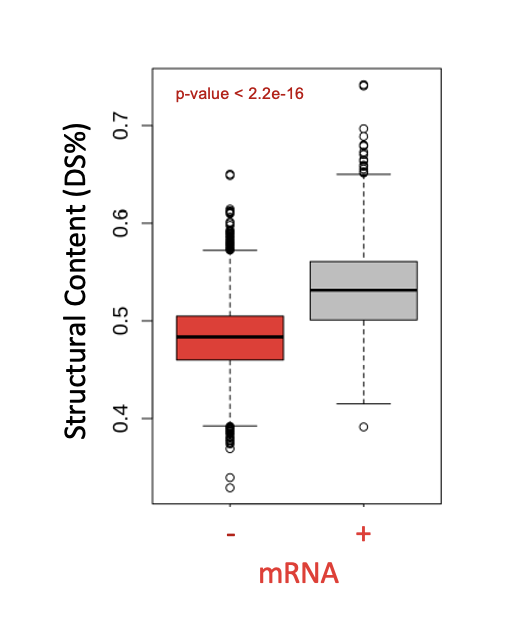

**Supplementary Figure 1.** Boxplot showing the distribution of structural content (i.e. DS% or double-stranded nucleotides) predicted in absence (“-” in red) and presence of ADAR (“+” in gray) for human mRNAs (22’000 RNAs).

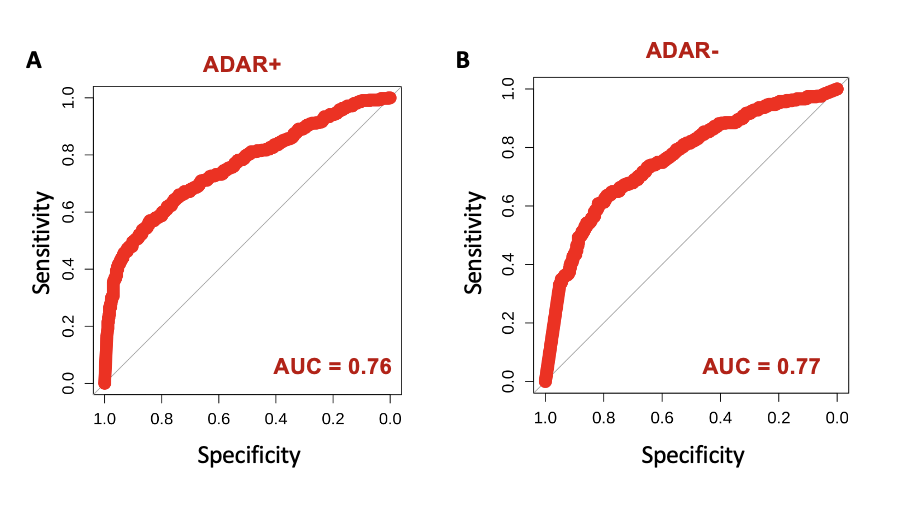

**Supplementary Figure 2.** Comparison between CROSSalive predictions and nextPARS experiments on NORAD. **A.** Performance of the CROSSalive method (ADAR+) on the 10% most structured regions (540 nucleotides) and the 10% least structured regions (540 nucleotides). The Area under the ROC curve is 0.76; **B.** Performance of the CROSSalive method (ADAR-) on the 10% most structured regions (540 nucleotides) and the 10% least structured regions (540 nucleotides). The Area under the ROC curve is 0.77.

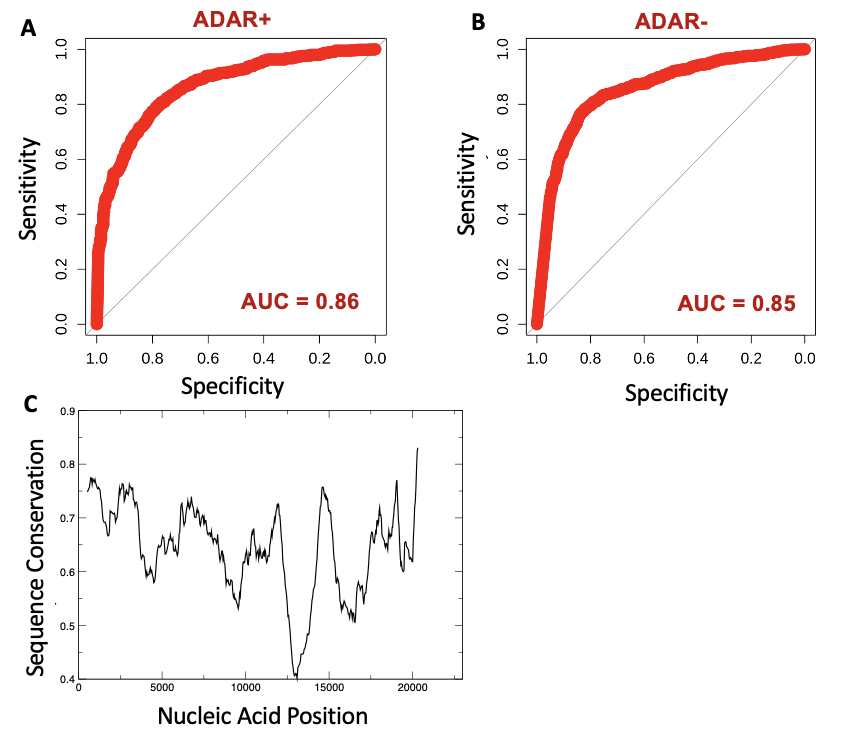

**Supplementary Figure 3.** Comparison between CROSSalive predictions and PARS experiments on NEAT and sequence conservation. **A.** Performance of the CROSSalive method (ADAR-). The 10% most structured regions (2200 nucleotides) and the 10% least structured regions (2200 nucleotides) are used. The Area under the ROC curve is 0.85. **B.** Performance of the CROSSalive method (ADAR-). The 10% most structured regions (2200 nucleotides) and the 10% least structured regions (2200 nucleotides). The Area under the ROC curve is 0.85; **C.** Sequence alignment of mouse and human NEAT indicates regions of high conservation.
